# Temporally feathered intensity modulated radiation therapy: a planning technique to reduce normal tissue toxicity

**DOI:** 10.1101/184044

**Authors:** Juan Carlos López Alfonso, Shireen Parsai, Nikhil Joshi, Andrew Godley, Chirag Shah, Shlomo A. Koyfman, Jimmy J. Caudell, Clifton D. Fuller, Heiko Enderling, Jacob G. Scott

## Abstract

**Purpose:** Intensity modulated radiation therapy (IMRT) has allowed optimization of three-dimensional spatial radiation dose distributions permitting target coverage while reducing normal tissue toxicity. However, radiation-induced normal tissue toxicity is a major contributor to patients’ quality of life and often a dose-limiting factor in the definitive treatment of cancer with radiation therapy. We propose the next logical step in the evolution of IMRT using canonical radiobiological principles, optimizing the temporal dimension through which radiation therapy is delivered to further reduce radiation-induced toxicity by increased time for normal tissue recovery. We term this novel treatment planning strategy “temporally feathered radiotherapy” (TFRT).

**Methods:** TFRT plans were generated as a composite of five simulated treatment plans each with altered constraints on particular hypothetical organs at risk (OARs) to be delivered sequentially. For each of these TFRT plans, OARs chosen for feathering receive higher doses while the remaining OARs receive lower doses than the standard fractional dose delivered in a conventional fractionated IMRT plan. Each TFRT plan is delivered a specific weekday, which in effect leads to a higher dose once weekly followed by four lower fractional doses to each temporally feathered OAR. We compared normal tissue toxicity between TFRT and conventional fractionated IMRT plans by using a dynamical mathematical model to describe radiation-induced tissue damage and repair over time.

**Results:** Model-based simulations of TFRT demonstrated potential for reduced normal tissue toxicity compared to conventionally planned IMRT. The sequencing of high and low fractional doses delivered to OARs by TFRT plans suggested increased normal tissue recovery, and hence less overall radiation-induced toxicity, despite higher total doses delivered to OARs compared to conventional fractionated IMRT plans. The magnitude of toxicity reduction by TFRT planning was found to depend on the corresponding standard fractional dose of IMRT and organ-specific recovery rate of sublethal radiation-induced damage.

**Conclusions:** TFRT is a novel technique for treatment planning and optimization of therapeutic radiotherapy that considers the non-linear aspects of tissue repair to optimize toxicity profiles. Model-based simulations of TFRT to carefully conceptualized clinical cases have demonstrated potential for normal tissue toxicity reduction in a previously described dynamical model of normal tissue complication probability (NTCP).

## Introduction

Since the advent of therapeutic radiotherapy, our understanding of radiation planning, delivery and effects have significantly grown. In parallel, the utilization of radiotherapy has increased, and currently it is estimated that about half of cancer patients benefit from curative or palliative therapy at some point during the course of their disease.^1^ The principle challenge in delivering safe, yet effective radiotherapy has been the balance of tumor control probability (TCP) against normal tissue complication probability (NTCP), termed the therapeutic ratio.^2^ The so called “widening” of the therapeutic ratio has been subject to research since the early implementation of radiotherapy. Specifically, treatment planning techniques have evolved from 2D to 3D planning, and now intensity modulated radiation therapy (IMRT), which has allowed optimization of physical dose distributions and avoidance of organs at risk.^3–5^ Moreover, clinicians can now also alter dose to the target as well, such as in heterogeneously dosing across a tumor volume, promoting dose intensification.^6^ This is all done while abiding by conventional dose constraints to normal tissues to qualify plan safety before patient delivery.^2^ The ability to spare organs and healthy tissues has been pivotal in improving quality of life during and following radiotherapy.^7^

Through these advancements, the four pillars of radiobiology have guided the understanding of radiation effect on tissue: *(i)* repair of sublethal damage, *(ii)* reassortment of cells within the cell cycle, *(iii)* repopulation, and *(iv)* reoxygenation. We accept sublethal damage repair and repopulation as the main drivers of dose limiting acute toxicity and hence fractionated radiotherapy still predominates in the clinic. The consistent fractional dose of radiation administered daily, delivers consistent insult to tumor cells while allowing time for normal tissue recovery between treatment fractions.^8^ Yet clinically, not all radiation-induced damage to organs at risk (OARs) is recovered with this interfractional interval. Taking the example of head and neck malignancies, treated to 70 Gy in 35 fractions, acute toxicities often still manifest midway or toward the end of most treatment courses. Despite reversibility of acute toxicity, when severe, it can lead to treatment breaks and compromise tumor control.^7^ Additionally, well-documented late effects contribute to the morbidity of radiation therapy, as these are often irreversible. The repopulation of normal tissues following peak acute toxicity can take weeks to months. We herein introduce a novel treatment planning strategy with the potential to reduce acute and late normal tissue toxicity.

Understanding basic radiobiology principles discussed above, we have developed a novel technique of optimizing radiation dose and fractionation, leveraging time to maximize normal tissue recovery, and therefore decrease toxicity without altering tumor dose. We hypothesize that if an organ at risk receives a once weekly higher than standard fractional dose of IMRT followed by lower fractional doses, the one week interval between the higher fractional doses will allow increased sublethal damage repair and repopulation. The novelty is in balancing normal tissue repair against radiation-induced damage using a non-linear model of dynamic NTCP. The focus of this study is introducing the theoretical radiation biology behind the advance. The proposed dynamical model of NTCP with a recovery term of normal tissue damage allows for the conceptual presentation of TFRT, and development of a predictor of TFRT benefit over conventionally planned IMRT. Possible clinical implications of reduced toxicity include improvement the quality of life issues, as well as potential for dose intensification to the tumor with similar toxicity profiles.

## Methods

### Temporally Feathered Radiation Therapy

We present a novel treatment planning strategy, which we term temporally feathered radiation therapy (TFRT), in which the fractional radiation dose delivered to OARs is altered to allow for increased normal tissue recovery of radiation-induced damage with respect to conventionally fractionated IMRT. A TFRT plan is generated as a composite of several iso-curative (i.e. same tumor dose) plans each with altered constraints on particular OARs of interest. In each of these TFRT plans, a single OAR would be deprioritized, allowing the optimization algorithm to reduce radiation dose and thereby toxicity to all other OARs. In practice, let us assume a planning target volume (PTV) with five surrounding OARs of interest prescribed a standard dose of 70 Gy in 35 fractions, similar to that commonly implemented for head and neck cancers.^7^ Further, let us consider that five treatment plans are developed, wherein each of the five OARs receives a relatively high fractional dose (d_H_) compared to the standard fractional dose (d_S_) once weekly, i.e. 2.0 Gy. A relatively lower (d_L_) fractional dose is then delivered the remaining four days of the week (see Figure 1). With this treatment planning strategy, though greater radiation-induced damage is induced by dH once weekly, it is offset by the lower fractional dose, d_L_, delivered over a greater amount of time, i.e. during the remaining four days. We then compare the composite of d_H_ and d_L_ to the corresponding standard fractional dose dS delivered to each OAR in a conventionally fractionated IMRT plan. In this hypothetical case, the TFRT plan is composed by 35 fractions, and each OAR of interest will receive 28 fractions of 0 ~ d_L_ ~ d_S_ and 7 fractions of d_H_ > d_S_ > 0. We consider that fractional doses d_L_ and d_H_ remain unaltered during the course of treatments. For demonstrative purpose, we focus on radiotherapy treatment plans which feather five OARs, though any number of OARs can be chosen for temporally feathering.

**Figure 1.**
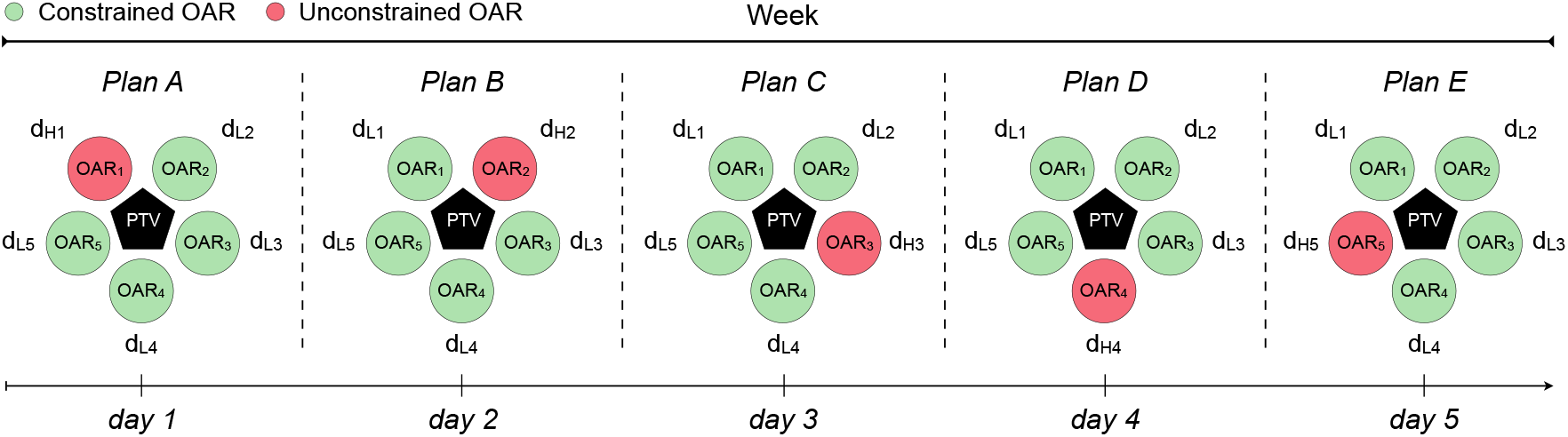
Schematic representation of treatment planning in temporally feathered radiation therapy (TFRT). The planning target volume (PTV) is in close proximity to five organs at risk (OARs). Each OAR *i* receives a higher fractional dose once weekly (d_Hi_), followed by lower fractional doses for the remaining four weekdays (d_Li_), *1 ≤ i ≤ 5*. The PTV is represented as a black pentagon, and circles represent the surrounding OARs. Green: constrained OAR. Red: unconstrained OAR.

### Biologically effective dose (BED) model

The Linear-Quadratic (LQ) model is currently the most widely used dose-response formulation in radiotherapy.^8–9^ The LQ model fit to *in vitro* cell survival experiments and incorporates the linear-quadratic behavior of observed cell survival curves.^9^ The linear component accounts for cell killing by DNA double strand breaks (DSBs) due to a single hit of radiation, whereas the quadratic component represents the lethal effects of two separate ionizing events that eventually cause DSBs.^8–10^ The surviving fraction (SF) of cells after *n* fractions of a radiation dose *d* is given by

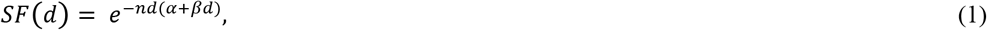

where α (Gy^−1^) and β (Gy^−2^) are tissue-dependent radiosensitivity parameters. It follows directly from the LQ-model that the biological effect (E) of *n* equally sized fractions of dose *d* is given by E = *nd* (*α* + *βd*). This equation can be manipulated to derive biologically effective dose (BED) calculations, which is a standard quantity allowing comparison of various radiotherapy fractionation schemes. BED is dependent on inherent biologic radiosensitivity of tissues, which is termed as the α to β ratio, α/β. This is also borrowed from the LQ model.^9–11^ The biologically effective dose^11–12^ is given by

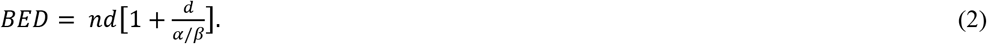

The BED equation above applies to conventionally fractionated radiation plans in which a same fractional dose (i.e. standard dose) is daily delivered. The BED for a standard daily treatment fraction (BED_S_) is given by

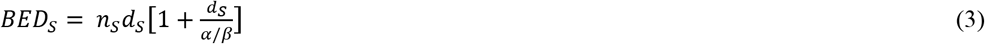

where n_S_ is the number of treatment fractions and d_S_ is the radiation dose per fraction. The BED of temporally feathered plans BEDTF is defined as follows

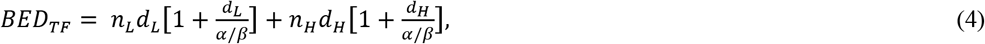

where n_L_ and n_H_ refer to the number of lower dose (d_L_) fractions and number of higher dose (d_H_) fractions, respectively. Lower-dose fractions deliver a radiation dose less than what would be delivered in a conventionally fractionated IMRT plan, 0 < d_L_ < d_S_. Similarly, higher-dose fractions deliver a radiation dose higher than what would be delivered in a conventionally fractionated IMRT plan, d_H_ > d_S_ > 0. We consider that fractional doses d_L_ and d_H_ remains unaltered during the course of treatments, and are homogeneously distributed on each OAR. The total number of fractions and their time of delivery remains the same in conventionally fractionated IMRT and TFRT plans, i.e. n_S_= n_L_ + n_H_, and we are not changing tumor dose, only dose to OARs.

### BED-based comparison of treatment plans

The difference in the BED delivered by a conventionally fractionated IMRT plan (S) of standard dose d_S_ and a temporally feathered (TF) radiation therapy plan is defined as ΔBED = BED_S_ – BED_TF_, see Equations (3) and (4).

### Dynamical model of normal tissue complication probability (NTCP)

We use a non-spatial dynamical mathematical model to simulate normal tissue response to radiation. This is a form of NTCP modeling, which is a quantitative measure of radiation-induced detriment to normal tissues^13–17^. The model is formulated as a logistic differential equation that describes the recovery of normal tissues (*N*) from sublethal radiation-induced damage given by

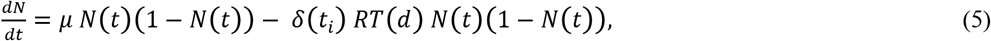

where the organ-specific parameter μ > 0 represents the recovery rate of radiation-induced damage. We consider the case that before radiation, the OAR is at tissue homeostasis with a 1% turnover rate, N(0) = 0.99. Then N(t) < N(0) represents the level of normal tissue damage by radiotherapy (small values of N(t) relate to severe damage), and (N(0) - N(t)) is used as an indication of the radiation-induced toxicity. The logistic differential Equation (5) used to model normal tissue recovery simulates a decay of toxicity to zero overtime. This is based on clinical observations revealing that not all patients develop late toxicities, and more importantly, that acute toxicities normally do go to zero on rather short time scales. Further, this model is used to compare conventionally fractionated and temporally fathered radiotherapy plans under the same conditions, which does not influence the ability to compare planning techniques.

The effect of radiation is included by the loss term *δ*(*t_i_*) *RT*(*d*) *N*(*t*)(1 − *N*(*t*)) in Equation (5), where *δ*(*t_i_*) is the Dirac-delta function equals to one at the time of irradiation *t_i_*, and zero otherwise. The structure of this loss term models the growing effect of radiation therapy with increasing number of treatment fractions. In fact, it is known that as treatment fractions accumulate the observed radiation-induced acute toxicities become increasingly apparent.^18^ Clinically, we observe normal tissue toxicities that increase in severity mid-way and towards the end of treatment. The function *RT*(*d*)= (1 − *e*^*−αd −βd*^2^^) is based on the radiobiological LQ-model in Equation (1). More precisely, *RT*(*d*) represents the “injured fraction” of normal cells receiving a radiation dose d, i.e. 1 – surviving fraction of cells. Thus, for low radiation doses the injured fraction of normal cells due to radiation must be small, thereby *RT*(*d*) must be close to zero. On the other hand, high radiation doses will result in more killed normal cells for which *RT*(*d*) tends to one. Further, we assume that both the delivery of each treatment fraction and response to radiation are instantaneous. We notice that similar dynamical models have been previously proposed to simulate the effect of radiation on brain^19–21^ and lung tumors^22^, as well as to define an organ-specific NTCP model^23^.

### NTCP-based comparison of treatment plans

We denote by ΔNTCP = N_S_(t_end_) - N_TF_(t_end_), the difference between OAR toxicity induced by a conventionally fractionated IMRT plan (NS(tend)) and a temporally feathered radiation therapy plan (N_TF_(t_end_)) at the end of treatment t_end_, see Equation (5). This means that positive values (ΔNTCP > 0) favor TFRT over IMRT plans.

### Overall and maximum potential benefit of TFRT over conventionally fractionated IMRT

We estimate the normal tissue toxicity reduction of TFRT over conventionally planned IMRT by using a term referred to as *overall potential benefit* (OPB_TF_). For any given combination of the organ-specific recovery rate *μ* and the fractional radiation dose d_S_ delivered by a conventionally fractionated IMRT plan, OPB_tf_ is the ratio of simulated TFRT plans with 0 < d_m_ ≤ d_L_ ≤ d_S_ and 0 < d_S_ ≤ d_H_ ≤ d_M_ that result in ΔNTCP > 0 and deliver higher total doses than the corresponding IMRT plans. d_m_ and d_M_ are the minimum lower dose (d_L_) and the maximum higher dose (d_H_) considered to generate the TFRT plans.

The *maximum potential benefit* (MAX_TF_) of TFRT over conventionally planned IMRT is the maximum ΔNTCP > 0 of simulated TFRT plans delivering higher total doses than the corresponding IMRT plans.

## Results

### BED model simulations

We first consider the BED model to compare TFRT and conventionally fractionated IMRT under varying conditions. To that end, we considered an OAR at a physiologic equilibrium and characterized by a α/β ratio of 3 Gy. Further, we simulated TFRT plans with d_m_ ≤ d_L_ ≤ d_S_ and d_S_ ≤ d_H_ ≤ d_M_ consisting of 28 fractions (n_L_) of d_L_ ≤ d_S_ and 7 fractions (n_H_) of d_H_> d_S_, and the corresponding conventionally fractionated IMRT plans delivering d_S_ in 35 fractions. For illustrative purposes, we have chosen d_m_= (d_S_ – 0.5 Gy) and d_M_= (d_S_+ 2.5 Gy) with a dose increment of 0.01 Gy between d_m_ and d_S_, and between d_S_ and d_M_.

Figure 2 illustrates ΔBED = (BED_S_ - BED_TF_) between different TFRT and conventionally planned IMRT plans, see Equations (3) and (4). Irrespective of d_S_, TFRT plans result only in a lower BED when the total dose (28 d_L_+ 7 d_H_) delivered to the OAR of interest is less compared to the standard IMRT plan (35 d_S_). Further, combinations of d_L_ and d_H_ exist in which BEDTF > BEDS even when the total dose by TFRT plans is less than in the conventionally fractionated IMRT plan. These results hold irrespective of the α/β ratio of the OAR of interest (see Supplementary Figure 1). The BED formulation does not account for the effect of interfractional normal tissue recovery of radiation-induced damage, and therefore is not a suitable model to evaluate the potential benefit of TFRT. This highlights the need for models, which account for the dynamic of normal tissue recovery from radiation-induced damage between treatment fractions to assess the feasibility of TFRT.

**Figure 2.**
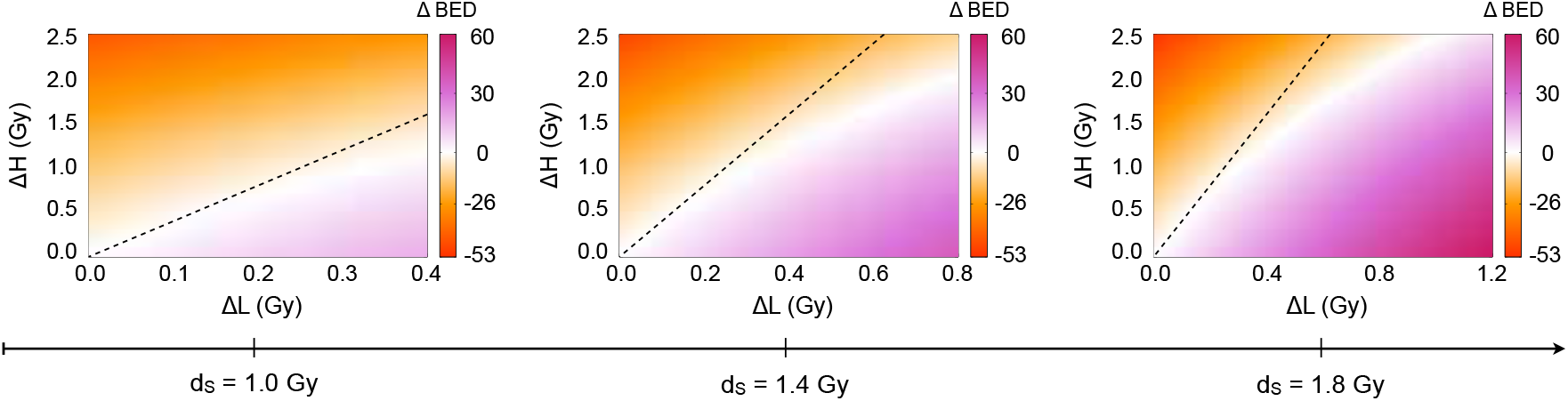
Comparison of conventionally fractionated IMRT and TFRT based on the biologically effective dose (BED) model. From left to right,ΔBED is represented in the divergent colormap for increasing doses d_S_. The x- and y-axes represent ΔL = d_S_ – d_L_ and ΔH = d_H_ – d_S_, respectively. The regions below and above the dashed lines represent combinations of d_L_ and d_H_ in which TFRT plans deliver lower and higher total doses compared to the corresponding IMRT plan delivering a fractional dose d_S_, respectively.

### Dynamical NTCP model simulations

We now simulate normal tissue complication probability (NTCP) of TFRT compared to conventionally fractionated IMRT implementing the dynamical NTCP model presented in Equation (5). As above, we consider an OAR with α/β= 3 Gy and a conventionally fractionated IMRT plan delivering a standard fractional dose d_S_ in 35 fractions. TFRT plans consist of 28 fractions (n_L_) of d_L_ ≤ d_S_ and 7 fractions (n_H_) of d_H_ > d_S_, with d_L_ and d_H_varying in the ranges [d_S_ - 0.5 Gy, d_S_] and [d_S_, d_S_+ 2.5 Gy], respectively. Model simulations reveal a range of treatment planning conditions in which TFRT plans reduce radiation-induced toxicity to OARs compared to conventional planned IMRT plans. These conditions are dependent on d_L_ and d_H_, as well as on the organ-specific recovery rates *μ*, associated with radiation-induced damage. This is shown in Figures 3 and 4, which represent ΔNTCP for TFRT and conventionally fractionated IMRT plans with varying *μ* and d_S_ values, respectively. We found that there exist combinations of d_L_ and d_H_ delivering higher total doses in TFRT plans as compared to conventionally fractionated IMRT plans (28 d_L_ + 7 d_H_ > 35 d_S_) but yet reduce the overall OAR toxicity. This is depicted by the regions above the dashed lines, but still in the beneficial (red: ΔNTCP > 0) regions, in the bottom panels of Figures 3 and 4.

**Figure 3.**
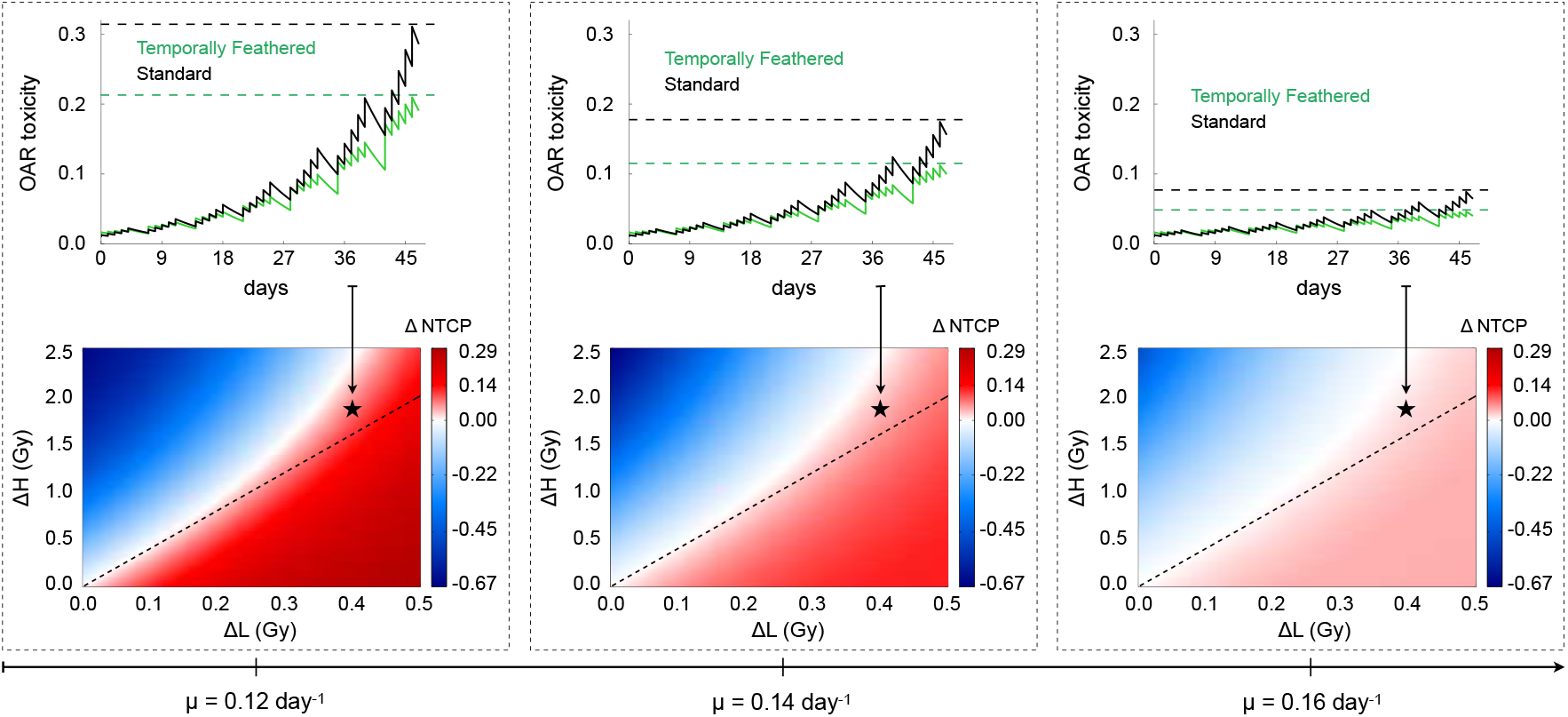
Comparison and representation of NTCP and OAR toxicity between conventionally fractionated IMRT and TFRT with varying organ-specific recovery rates (*μ*). Bottom panels, from left to right, ΔNTCP (color-bar: red is beneficial, blue is detrimental) are represented for d_S_= 1.2 Gy and increasing values of *μ*. The x- and y-axes represent ΔL = d_S_ – d_L_ and ΔH = d_H_ – d_S_, respectively. The regions below and above the dashed lines represent combinations of d_L_ and d_H_ in which TFRT plans deliver lower and higher total composite doses compared to the corresponding IMRT plan delivering a fractional dose d_S_, respectively. **Top panels** show the time-evolution of OAR toxicity induced by the IMRT (black) and TFRT (green) plans corresponding to the location marked by stars in bottom panels. Dashed lines represent the time points at which NTCP of IMRT and TFRT plans are compared.

**Figure 4.**
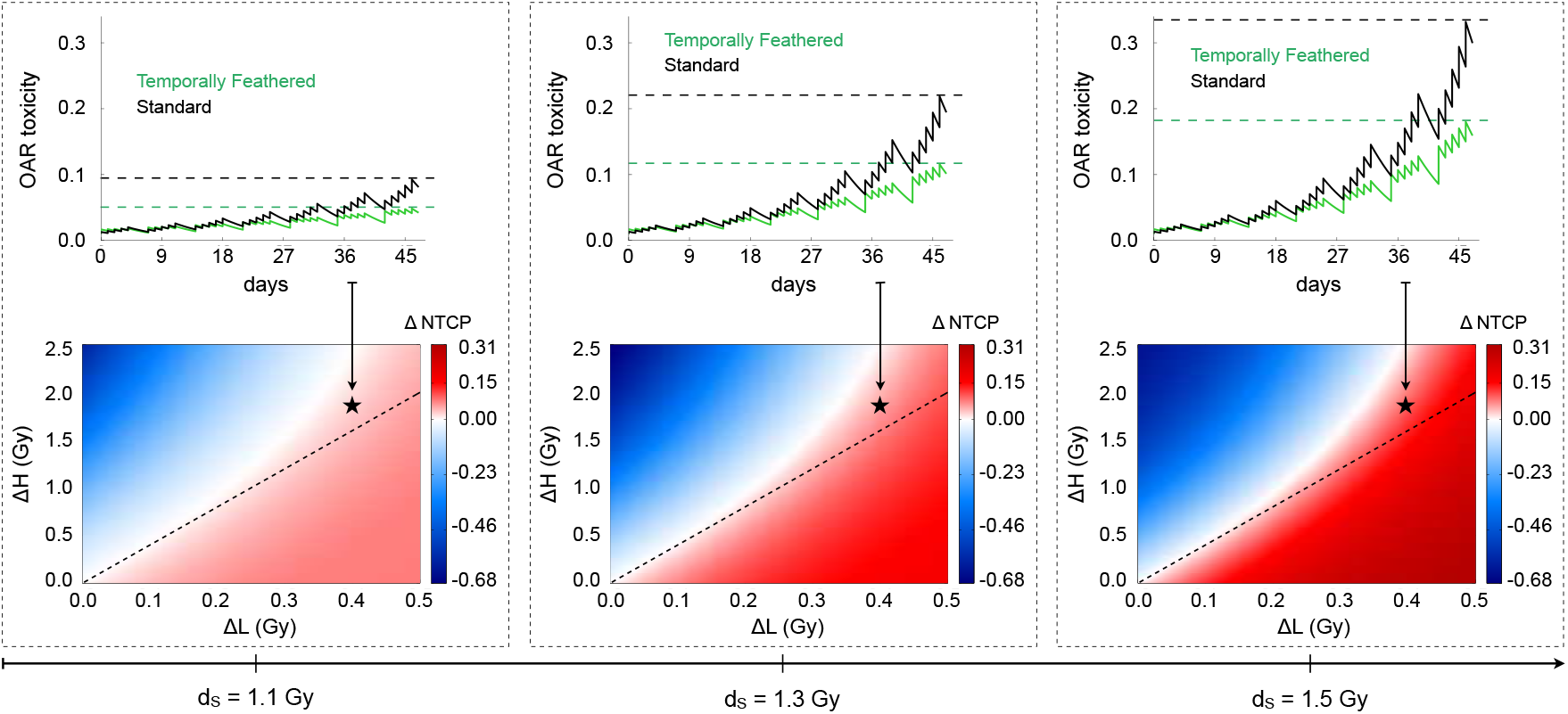
Comparison and representation of NTCP and OAR toxicity between conventionally fractionated IMRT and TFRT with varying standard fractional doses (dS). Bottom panels, from left to right, ΔNTCP (color-bar: red is beneficial, blue is detrimental) are represented for *μ*= 0.15 day^−1^ and increasing fractional doses d_S_ IMRT plans. The x- and y-axes represent ΔL = d_S_ – d_L_ and ΔH = d_H_ – d_S_, respectively. The regions below and above the dashed lines represent combinations of d_L_ and d_H_ in which TFRT plans deliver lower and higher total composite doses compared to the corresponding IMRT plan delivering a fractional dose d_S_, respectively. **Top panels** show the time-evolution of OAR toxicity induced by the IMRT (black) and TFRT (green) plans corresponding to the location marked by stars in bottom panels. Dashed lines represent the time points at which NTCP of IMRT and TFRT plans are compared.

The therapeutic gain by TFRT plans increases as the treatment progresses. This is shown in the top panels of Figures 3 and 4, where irrespective of the values of *μ* and standard fractional doses d_S_ considered, the difference in the radiation-induced OAR toxicity, i.e. N_S_(t) - N_TF_(t), see Equation (5), by conventionally fractionated IMRT and TFRT plans progressively increases with the number of treatment fractions. Furthermore, Figures 3 and 4 shown the difference between OAR toxicity induced by conventional planned IMRT and TFRT plans at the end of treatment (ΔNTCP) is greater with decreasing values of *μ* and increasing fractional doses d_S_. Thus, TFRT is more beneficial for reducing radiation-induced toxicity in OARs with low recovery rates *μ* and receiving high standard fractional doses d_S_ with conventional planned IMRT.

Figure 5 summarizes the impact of organ-specific and treatment parameters on the potential benefit of TFRT over conventionally fractionated IMRT. For each combination of *μ* and d_S_ considered, Figures 5(A) and 5(B) show the overall potential benefit (OPBTF) and maximum potential benefit (MAXTF) of TFRT plans with (d_S_ – 0.5 Gy) < d_L_ ≤ d_S_ and d_S_ ≤ d_H_ ≤ (d_S_+ 2.5 Gy) over the corresponding IMRT plans delivering a standard fractional dose d_S_. Figure 5 shows that while keeping constant d_S_ or *μ*, and varying the other parameter, the OPBTF and MAXTF of TFRT increase until a maximum level and then decrease again. This suggests that for each OAR characterized by a specific recovery rate *μ*, TFRT plans can be designed to reduce OAR toxicity if the standard fractional dose d_S_ delivered by a conventionally fractionated IMRT plans lies in a certain range. Furthermore, there exists an optimal dose d_S_ in that range for which OAR toxicity reduction with TFRT is greater. Similarly, OARs receiving a specific standard fractional dose d_S_ with conventional planned IMRT can be temporally feathered if they have a recovery rate *μ* is in a certain range. This evidences that both d_S_ and *μ* must be considered together when determining the OAR toxicity reduction from TFRT over IMRT. Supplementary Figures 2 and 3 show that OPB_TF_ and MAX_TF_ of TFRT over conventionally fractionated IMRT also depend on the specific α/β ratio of the OARs of interest. Those figures illustrate that, when applied on OARs characterized by different α/β ratios, TFRT continues to represent a theoretically valuable treatment planning strategy to reduce radiation-induced OAR toxicity.

**Figure 5.**
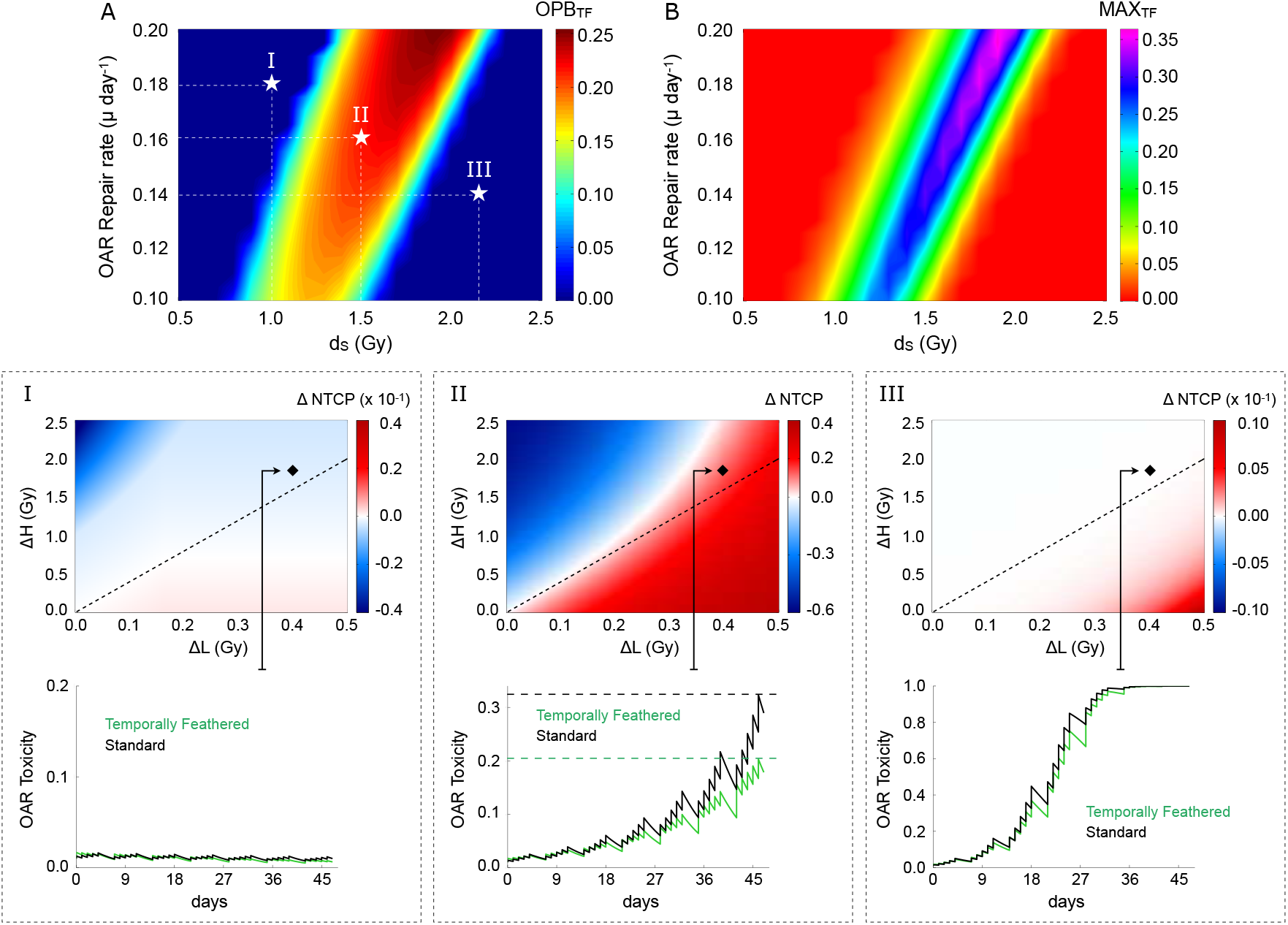
Comparison of conventionally fractionated IMRT and TFRT with respect to the standard fractional dose (d_S_) and organ-specific recovery rate (*μ*). **(A)** Overall potential benefit (OPB_TF_) and **(B)** maximum potential benefit (MAXTF) of TFRT over conventional planned IMRT. (**I-III**) **Top panels** represent the single cases marked by stars in (A). The x- and y-axes represent ΔL = d_S_ – d_L_ and ΔH = d_H_ – d_S_, respectively. **Bottom panels** show time-evolution of OAR toxicity induced by the IMRT (black) and TFRT (green) plans corresponding to the location marked by diamonds in the top panels.

## Discussion

Through the years, researchers in the field of radiation oncology and medical physics have been innovating new ways of widening the therapeutic ratio by either increasing tumor control probability (TCP) or decreasing normal tissue complication probability (NTCP). Recent works have shown the potential of spatiotemporal fractionation schemes delivering distinct radiation dose distributions in different fractions to improve the therapeutic ratio.^6,24-25^ The goal has been to maximize the mean BED in the tumor and to minimize the mean BED in normal tissues by hypofractionating parts of the tumor while delivering approximately identical doses to the surrounding normal tissue. This planning strategy has been shown to result in spatiotemporal fractionation treatments that can achieve substantial reductions in normal tissue dose. However, the effect of interfractional normal tissue recovery of radiation-induced damage has not been taken into account, which when considered could lead to further reduce treatment side effects. In this study, we consider the non-linear aspects of normal tissue repair to optimize toxicity profiles without compromising tumor control. We introduce TFRT as a novel treatment planning strategy that alters fractional radiation doses delivered to OARs over time with the hypothesis that this will lead to greater overall normal tissue recovery of radiation-induced damage through the course of treatment. This paper is an exercise in theory that TFRT has the potential to reduce normal tissue toxicity if the assumptions made in Equation (5) are valid, namely the recovery term. Conceptually, TFRT planning capitalizes on the nonlinearity of normal tissue recovery, allowing non-intuitively for more occasional sublethal damage repair and prolonged repopulation phases even in the face of higher total dose delivered at the end of treatment. For this purpose, we used the LQ-model to describe the immediate radiation response of normal tissue and a dynamical NTCP model to describe normal tissue repair of radiation-induced damage during fractions and over the entire treatment time.

The dynamical NTCP model considered in this work parallels prior similar models which have been proposed to simulate radiation effects on different tumor types^19–22^, as well as in healthy tissues^23^. The current LQ-model and BED formulation are limited by the lack of a temporal recovery term of radiation-induced damage, and therefore are not suitable models to compare radiation schema with various fractionations over time. Thus, a dynamical NTCP model with normal tissue recovery is pivotal to demonstrate potential toxicity reduction with TFRT *in silico*. The proposed NTCP model is sensitive to parameter changes including OAR recovery rate *μ*, α/β ratio and standard fractional dose d_S_. As these parameters represent phenomenological values, this model is personalizable to different clinical scenarios, which is relevant for recent work on Genomic Radiation Dosing revealing a wide heterogeneity in response to radiation therapy.^26^ Understanding the effect of TFRT on particular OARs allows for OAR prioritization to be used in optimization methods. Additional simulations under altered environments are included in the Supplementary Material. We emphasize that OAR-specific parameters may crucially determine the potential benefit of TFRT in decreasing normal tissue toxicity, which has important implications for clinical trial design. The concept of overall potential benefit is used to determine the potential of TFRT plans in reducing normal tissue toxicity as compared to conventionally fractionated IMRT plans (see Figure 5, and Supplementary Figures 2 and 3). Candidates for TFRT are patients with target volumes in close proximity to the organs at risk, in which conventionally planned IMRT leads to fractional doses to OARs near tolerance and with low recovery rates of radiation-induced damage.

While the focus of the work herein is to introduce the theoretical concept of TFRT, future work will need to evaluate feasibility of patient-specific TFRT plans with currently available treatment planning systems. Endpoints for TFRT plan dosimetry and safety evaluation should be defined including dose-volume histogram (DVH) based metrics.^27–29^ Before application in the clinic, an optimized workflow must be developed. The radiation plans must be developed in tandem, otherwise the creation, optimization, evaluation and quality assurance (QA) of five separate plans will not prove feasible in the clinic. Currently organ defined recovery rates (μ) and radiosensitivity parameters *α* and *β* values are not reliably defined, limiting the utility of this model at this time for clinical decision-making. However, after prospective clinical implementation and careful data gathering, efforts will be directed toward elucidating these parameters for various OARs. Based on radiobiological properties of the OARs involved, clinicians can be informed regarding the number of OARs to be feathered with optimized timing. In future work, temporal optimization will be combined with spatial optimization and evaluation of partial volumes of organs rather than their entirety. This has been described previously as different portions of the organs may preferentially contribute to toxicity.^29^ These concepts can be applied to any disease site in which the target is within close proximity to multiple surrounding organs at risk and are not only limited to head and neck malignancies.

## Conclusions

We introduce a novel strategy of treatment planning termed temporally feathered radiation therapy (TFRT), by which the radiation dose to organs at risk is optimized through time, which suggests an opportunity to improve normal tissue recovery from radiation-induced damage. *In silico* simulations using a dynamical NTCP model, accounting for normal tissue recovery demonstrate the potential of TFRT to reduce OAR toxicity compared to conventionally planned IMRT. Future work is focused on feasibility of TFRT planning using current treatment planning systems and ultimately translation to be prospectively evaluated in clinic.

## Acknowledgements

J. C. L. Alfonso gratefully acknowledges the funding support of the German Federal Ministry of Education and Research (BMBF) for the eMED project SYSIMIT (01ZX1308D). J. G. Scott thanks the NIH Loan repayment program for their generous support of his research. This research is partially supported by the Andrew Sabin Family Foundation; C. D. Fuller is a Sabin Family Foundation Fellow. C. D. Fuller receives direct funding and salary support from the National Institutes of Health (NIH), including: the National Institute for Dental and Craniofacial Research Award (1R01DE025248-01/R56DE025248-01); a National Science Foundation (NSF), Division of Mathematical Sciences, Joint NIH/NSF Initiative on Quantitative Approaches to Biomedical Big Data (QuBBD) Grant (NSF 1557679); the NIH Big Data to Knowledge (BD2K) Program of the National Cancer Institute (NCI) Early Stage Development of Technologies in Biomedical Computing, Informatics, and Big Data Science Award (1R01CA214825-01); NCI Early Phase Clinical Trials in Imaging and Image-Guided Interventions Program (1R01CA218148-01); an NIH/NCI Cancer Center Support Grant (CCSG) Pilot Research Program Award from the UT MD Anderson CCSG Radiation Oncology and Cancer Imaging Program (P30CA016672) and an NIH/NCI Head and Neck Specialized Programs of Research Excellence (SPORE) Developmental Research Program Award (P50 CA097007-10). C. D. Fuller receives salary support from the Patient-Centered Outcomes Research Institute (PCORI). C. D. Fuller has received direct industry grant support and travel funding from Elekta AB.

## Supplementary Material

**Supplementary Figure 1:**
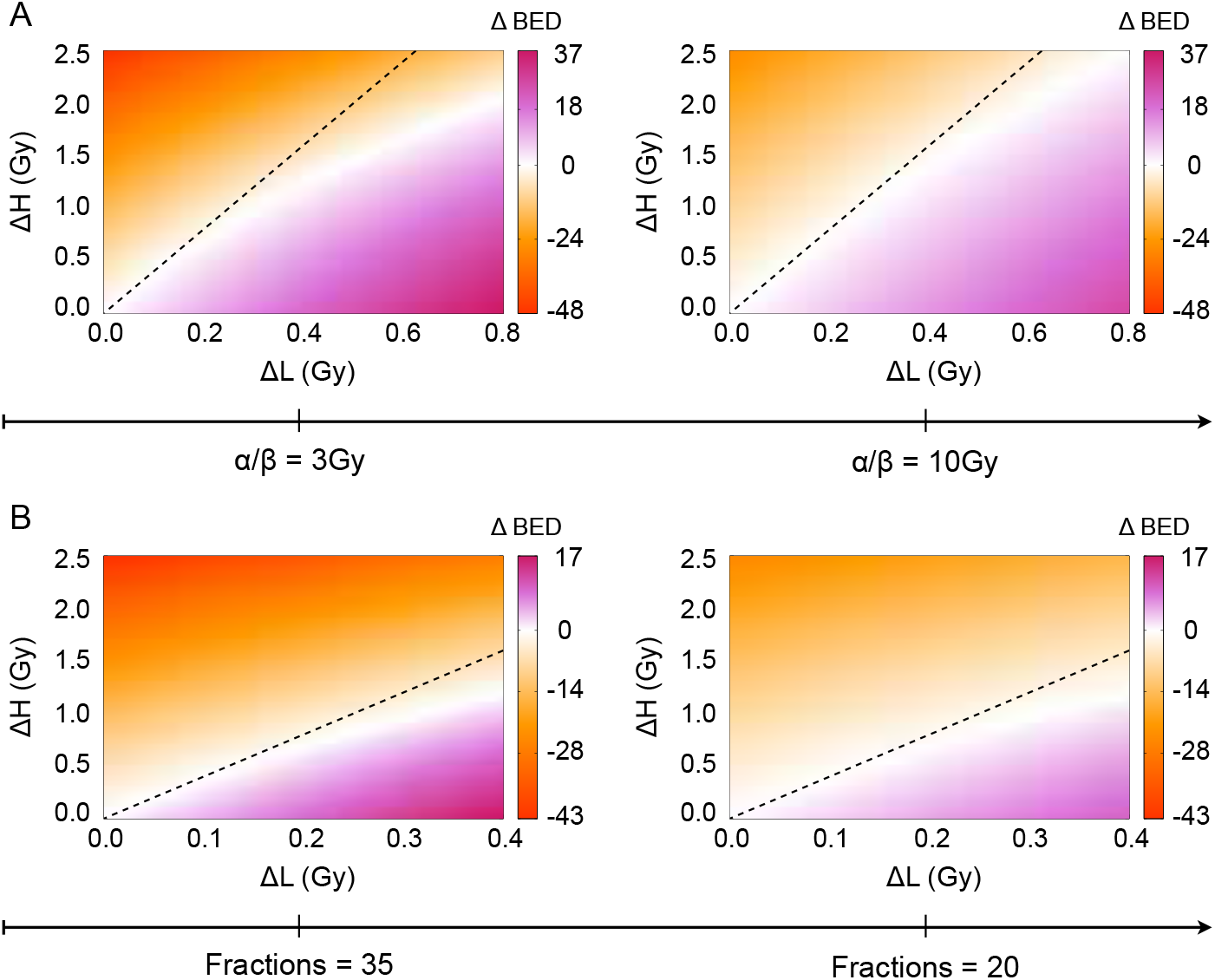
Comparison of conventionally fractionated IMRT and TFRT based on the biologically effective dose (BED) model. **(A)** From left to right, ΔBED is represented in the divergent colormap for different α/β ratios considering a conventional planned IMRT plan of d_S_= 1.4 Gy in 35 fractions, and TFRT plans of d_L_ < d_S_ in 28 fractions and d_H_> d_S_ in 7 fractions. **(B)** ΔBED is represented in the divergent colormap for an OAR with α/β = 3 Gy considering conventional planned IMRT plans of d_S_ = 1.0 Gy in 35 (left) and 20 (right) fractions, and TFRT plans of d_L_ ≤ d_S_ in 28 (left) and 16 (right) fractions and d_H_ > d_S_ in 7 (left) and 4 (right) fractions. The x- and y-axes represent ΔL = d_S_ – d_L_ and ΔH = d_H_ – d_S_, respectively. The regions below and above the dashed lines represent combinations of d_L_ and d_H_ in which TFRT plans deliver lower and higher total doses compared to the corresponding IMRT plan delivering a fractional dose d_S_, respectively.

**Supplementary Figure 2:**
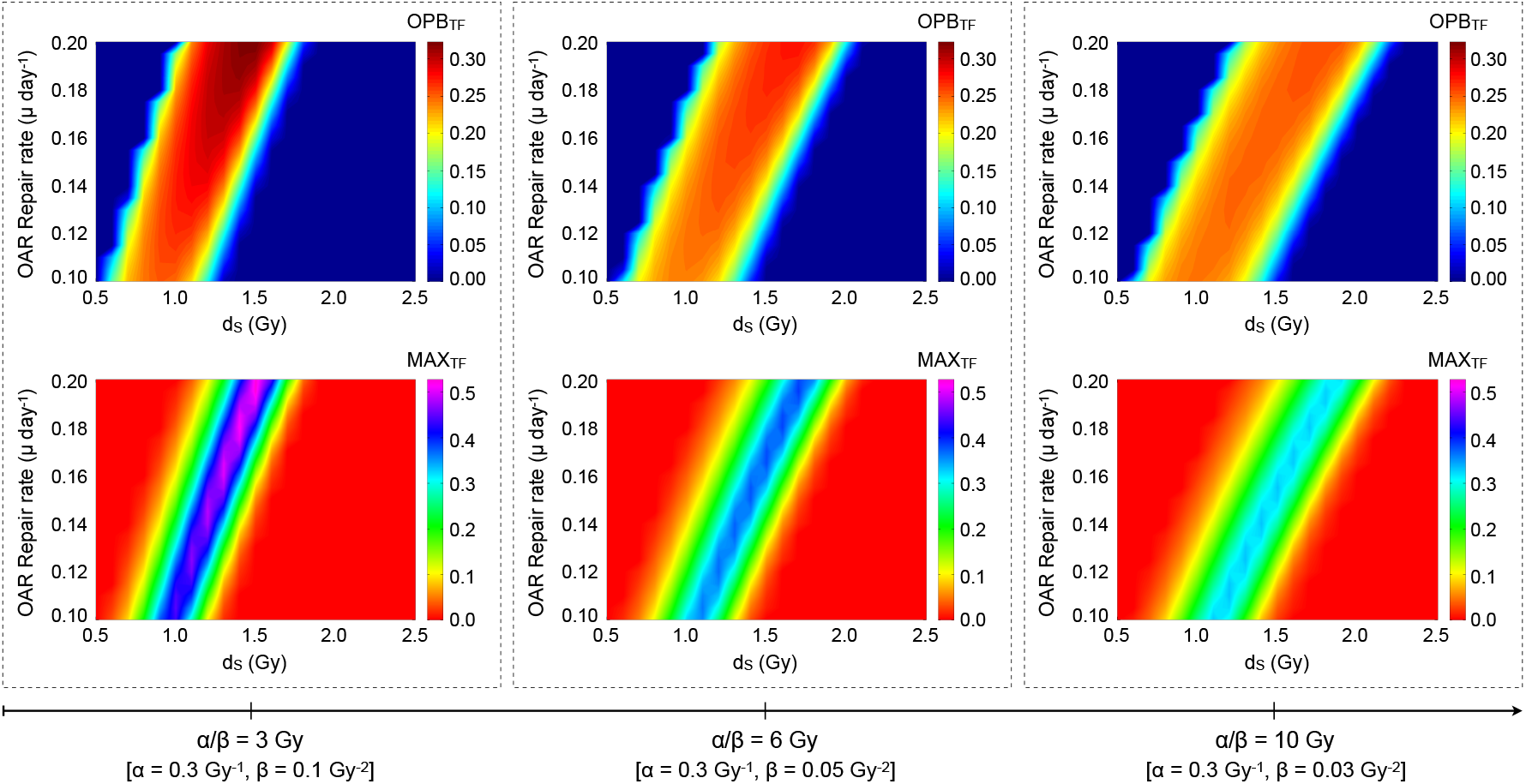
Comparison of conventionally fractionated IMRT and TFRT with respect to the standard fractional dose (d_S_) and organ-specific recovery rate *(μ*). **(Top panels)** Overall potential benefit (OPB_TF_) and **(Bottom panels)** maximum potential benefit (MAX_TF_) of TFRT over conventional planned IMRT for OARs of different *α/β* ratios with varying *β*. For each combination of *μ* and d_S_, ΔNTCP values are obtained as a result of comparing a conventional planned IMRT plan delivering d_S_ in 35 fractions, and TFRT plans consisting in 28 fractions of (d_S_ – 0.5 Gy) ≤ d_L_ ≤ d_S_ and 7 fractions of d_S_ ≤ d_H_ ≤ (d_S_ + 2.5 Gy).

**Supplementary Figure 3:**
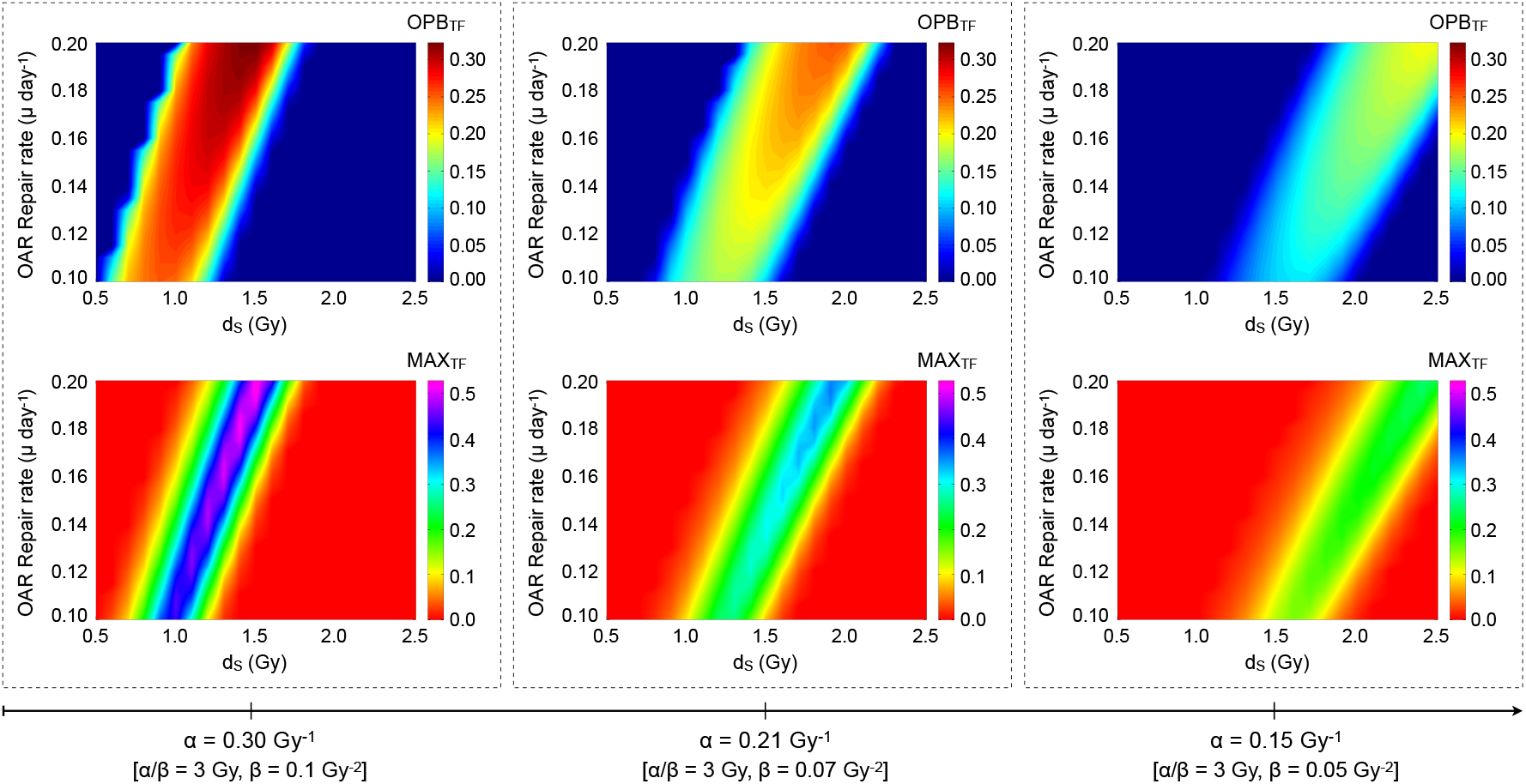
Comparison of conventionally fractionated IMRT and TFRT with respect to the standard fractional dose (d_S_) and organ-specific recovery rate (μ). **(Top panels)** Overall potential benefit (OPBTF) and **(Bottom panels)** maximum potential benefit (MAXTF) of TFRT over conventional planned IMRT for an OAR of *α/β*= 3 Gy with varying *α* and *β*. For each combination of μ and d_S_, ΔNTCP values are obtained as a result of comparing a conventional planned IMRT plan delivering d_S_ in 35 fractions, and TFRT plans consisting in 28 fractions of (d_S_ – 0.5 Gy) < d_L_ ≤ d_S_ and 7 fractions of d_S_ < dH < (dS + 2.5 Gy).

